# Differential Aging Effects on Implicit and Explicit Sensorimotor Learning

**DOI:** 10.1101/2024.07.02.601091

**Authors:** Elizabeth Cisneros, Sheer Karny, Richard B. Ivry, Jonathan S. Tsay

**Affiliations:** Department of Psychology, University of California, Berkeley; Helen Wills Neuroscience Institute, University of California, Berkeley; Department of Psychology, Carnegie Mellon University

## Abstract

Deterioration in motor control is a hallmark of aging, significantly contributing to a decline in quality of life. More controversial is the question of whether and how aging impacts sensorimotor learning. We hypothesized that the inconsistent picture observed in the current literature can be attributed to at least two factors. First, aging studies tend to be underpowered. Second, the learning assays used in these experiments tend to reflect, to varying degrees, the operation of multiple learning processes, making it difficult to make inferences across studies. We took a two-pronged approach to address these issues. We first performed a meta-analysis of the sensorimotor adaptation literature focusing on outcome measures that provide estimates of explicit and implicit components of adaptation. We then conducted two well-powered experiments to re-examine the effect of aging on sensorimotor adaptation, using behavioral tasks designed to isolate explicit and implicit processes. Convergently, both approaches revealed a striking dissociation: Older individuals exhibited a marked impairment in their ability to discover an explicit strategy to counteract a visuomotor perturbation. However, they exhibited enhanced implicit recalibration. We hypothesize that the effect of aging on explicit learning reflects an age-related decline in reasoning and problem solving, and the effect of aging on implicit learning reflects age-related changes in multisensory integration. Taken together, these findings deepen our understanding of the impact of aging on sensorimotor learning.

## Introduction

People are living longer with the majority of the population now surpassing the age of 60. The World Health Organization projects that by 2050, the population of people aged 60 years and older will double, rising from the current estimate of 1 billion to over 2 billion people (*Ageing and health*, 2018). As we age, there is a noticeable decline in mental and physical capabilities, though the rate of decline varies markedly between domains (Hedden & Gabrieli, 2004). In terms of physical capabilities, healthy aging is accompanied by a gradual deterioration of motor control (Contreras-Vidal et al., 1998; Darling et al., 1989; Diggles-Buckles & Vercruyssen, 1990; Seidler et al., 2002, 2010; Zapparoli et al., 2022), increasing the susceptibility to injuries, requiring more physical assistance, and diminishing the quality of life (Alexander et al., 1992; Reinkensmeyer & Patton, 2009; Sienko et al., 2008; Tinetti et al., 1988).

Age-related motor control deficits arise from a variety of structural changes in the neuromuscular system such as muscle atrophy (Naruse et al., 2023) and degeneration of sensory receptors (Johnsson & Hawkins, 1972). Other changes, such as the reduction in dopaminergic neurons which, in extreme cases, can result in Parkinson’s disease, are of more central origin. Motor control deficits in healthy aging may also be compounded by declines in processes associated with learning. In the present study, we focus on the effect of age on sensorimotor adaptation, the process of reducing motor errors through feedback and practice (Krakauer et al., 2019). A more aged sensorimotor system may not be as responsive to changes in the body (e.g., maintaining a constant force despite muscle fatigue) and environment (e.g., knitting with a new set of needles).

The effect of aging on motor adaptation remains controversial. Some studies show age-related declines in motor adaptation (Bock, 2005; Heuer & Hegele, 2008; Roller et al., 2002b) while others show comparable (Buch et al., 2003; Vachon et al., 2020) or even a superior (Seidler, 2006; Vandevoorde & Orban de Xivry, 2019) capacity for adaptation in older compared to younger adults. Two methodological factors may contribute to this inconsistency. First, most studies in this area are underpowered, involving modest sample sizes drawn from a limited age range (Roller et al., 2002a; Seidler, 2006). As such, the results may have low reliability and sensitivity.

Second, the learning assays used in these experiments tend to reflect the operation of multiple learning processes. Research over the past decade has highlighted how performance changes in sensorimotor adaptation tasks can reflect the joint operation of implicit and explicit learning processes. Implicit recalibration keeps movements coordinated in an automatic and subconscious manner, while explicit strategies flexibly correct for motor errors in a deliberate and conscious manner (for reviews, see (Huberdeau et al., 2015; Kim et al., 2020)). Considering that the relative contribution of these processes varies with the experimental context and dependent variable used to asses learning (Maresch et al., 2021, 2020; Ranganathan et al., 2021), the mixed findings in the literature may stem from differential effects of aging on distinct learning processes.

## Results

### Meta Analysis: Differential Aging Effects on Implicit and Explicit Sensorimotor Learning

#### Overview of studies included in the meta-analysis

As a first approach to ask how healthy aging impacts motor adaptation, we conducted a meta-analysis that encompassed studies that compared sensorimotor adaptation in younger and older adults. We focused on two measures of adaptation. First, we looked at performance after prolonged exposure to a perturbation, or what we will refer to as late adaptation. This phase includes contributions from both implicit and explicit processes. As such, this measure serves as an indicator of overall adaptation or task success. Second, we looked at performance once the perturbation has been removed. The deviation in heading angle away from the target in this post-learning phase (dubbed an “aftereffect”) provides a targeted measure of implicit recalibration. After applying a series of inclusion and exclusion criteria (Figure 1, see Methods), we identified 55 datasets across 44 studies that met our inclusion criteria. Note that some studies encompass multiple experiments with varying experimental parameters, thereby accounting for the presence of more datasets than studies. Reflective of the interest in this question, papers meeting our inclusion criteria span research conducted over the last three decades (Figure 2A) and collectively involving a total of 2326 participants.

**Figure 1.**
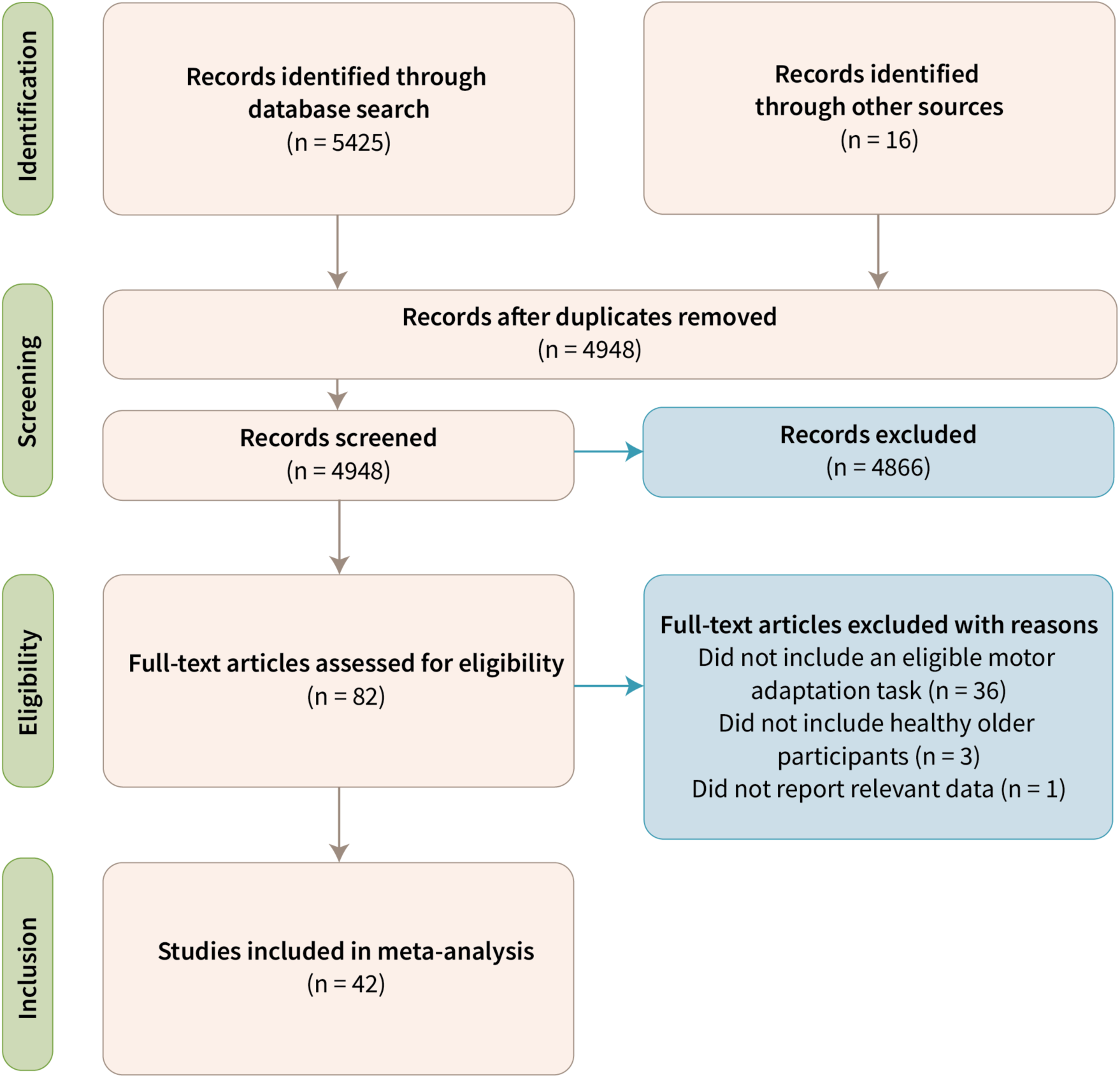
PRISMA flow diagram. A total of 44 studies (55 datasets) met our eligibility and inclusion criteria.

**Figure 2.**
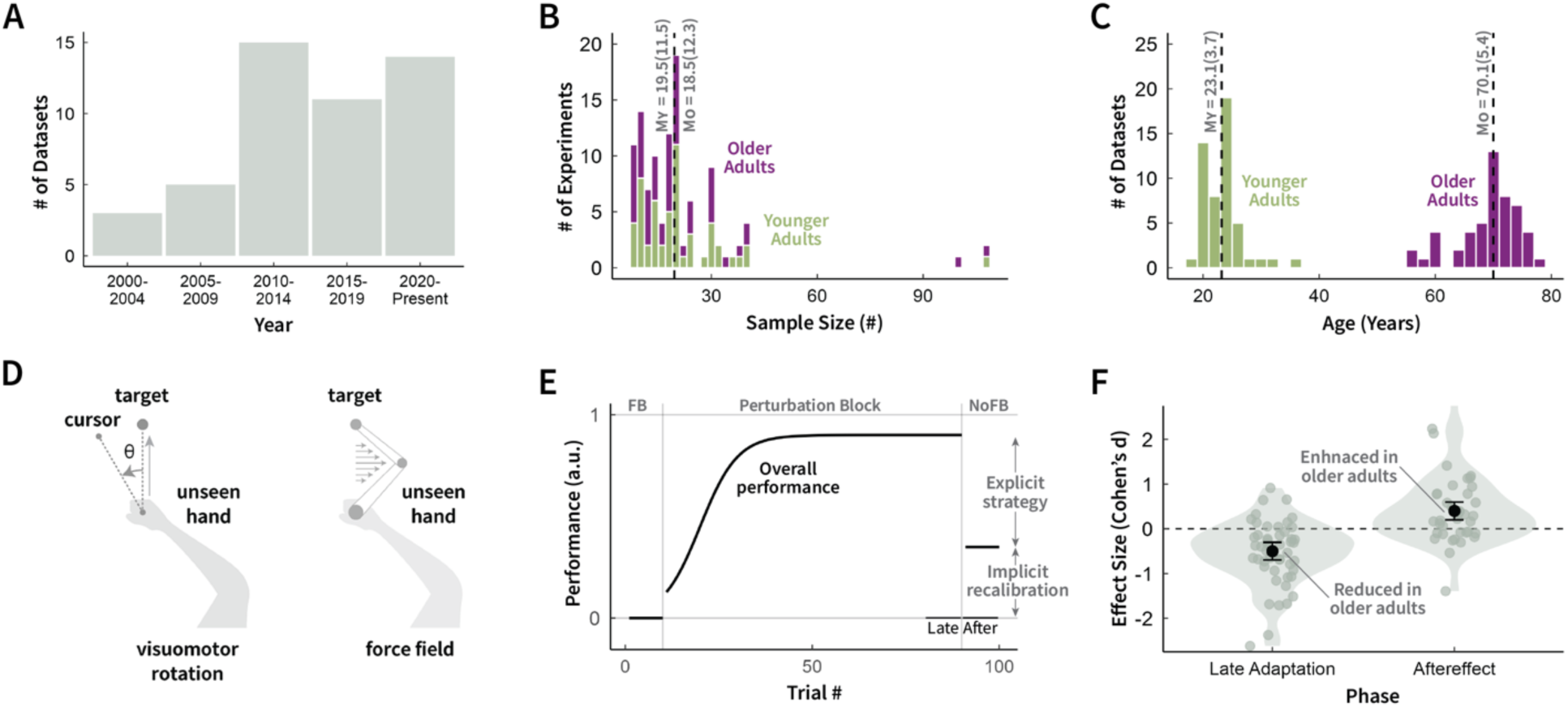
Overview of meta-analysis examining the impact of age on motor adaptation. **A-C)** Histograms indicating **A)** the number of datasets spanning 2000 and 2023, **B)** the sample size per dataset for younger (green) and older adults (purple), and **C)** the average age of younger and older adults per dataset. Dashed line represents the median (IQR). **D)** Schematics illustrating visuomotor rotation (left) and force-field (right) adaptation tasks (see Methods). **E)** Schematized learning curve with baseline, perturbed feedback, and aftereffect (no feedback) phases. Late adaptation during the perturbation block measures the total extent of learning, reflecting the contribution of both implicit and explicit learning processes. During the no-feedback aftereffect block, the perturbation is removed, and the participant is instructed to abandon their explicit strategy and reach straight to the target. In this way, the aftereffect is designed to provide a measure of implicit recalibration. **F)** Meta-analysis results. Effect sizes (Cohen’s d) are shown for both late adaptation and aftereffect measures. Positive effect sizes denote an enhancement in older adults; negative effect sizes denote a reduction in older adults. Individual translucent dots represent individual studies. The black dot denotes the overall effect size, and error bars denote the 95% confidence interval.

We did not find evidence of a publication bias for measures related to either late adaptation (X^2^(1) = 1.1, p = 0.3) or the aftereffect (X^2^(1) = 1.8, p = 0.2). However, we did find that most studies in the literature recruited small sample sizes and were likely to be statistically underpowered to detect age-related effects in motor adaptation (Figure 2B; Median N per group [IQR], Younger: 19.5 [13.3, 24.8]; Older: 18.5 [12, 24.3]; Median Statistical Power [IQR]: 0.4 [0.1, 0.8], with α = 0.05). There were two studies with relatively larger sample sizes (Li et al., 2021; Wolpe et al., 2020).

#### The detrimental effect of aging on motor adaptation

Fifty datasets compared late adaptation between older and younger adults (Age Mean ± SD, Younger: 23.3 ± 3.3; Older: 68.9 ± 5.4 years old (Figure 2E). The impact of aging on late adaptation was heterogenous (I^2^ = 73%, p < 0.001), with effect sizes ranging between -2.6 (a negative effect denotes reduced late adaptation in older adults) and 0.9 (a positive effect denotes enhanced late adaptation in older adults). However, when considering the aggregate data, we observed a detrimental effect of aging on motor adaptation: Late adaptation was significantly *reduced* in older compared to younger adults (Figure 2F & Figure 3; d = -0.5; 95% CI [-0.7, -0.3]; t(49) = -5.31; p < 0.001). This age-related decline was even more pronounced when the meta-analysis was restricted to only those that were adequately powered (power > 0.8; d = -1.4; 95% CI [-1.8, -1.0]; t(11) = -7.15; p < 0.001). Interestingly, the age-related decline in the extent of late adaptation did not differ between experimental tasks (visuomotor vs force-field adaptation: Q = 2.5, p = 0.14), perturbation sizes (range of visuomotor rotations: [1.8° - 90°]; β = -0.02, p = 0.3), or the number of visual targets (range: [1 - 10]; β = 0.08, p = 0.2). Thus, across a broad range of contexts, there is an age-related decline in overall performance on sensorimotor adaptation tasks.

**Figure 3:**
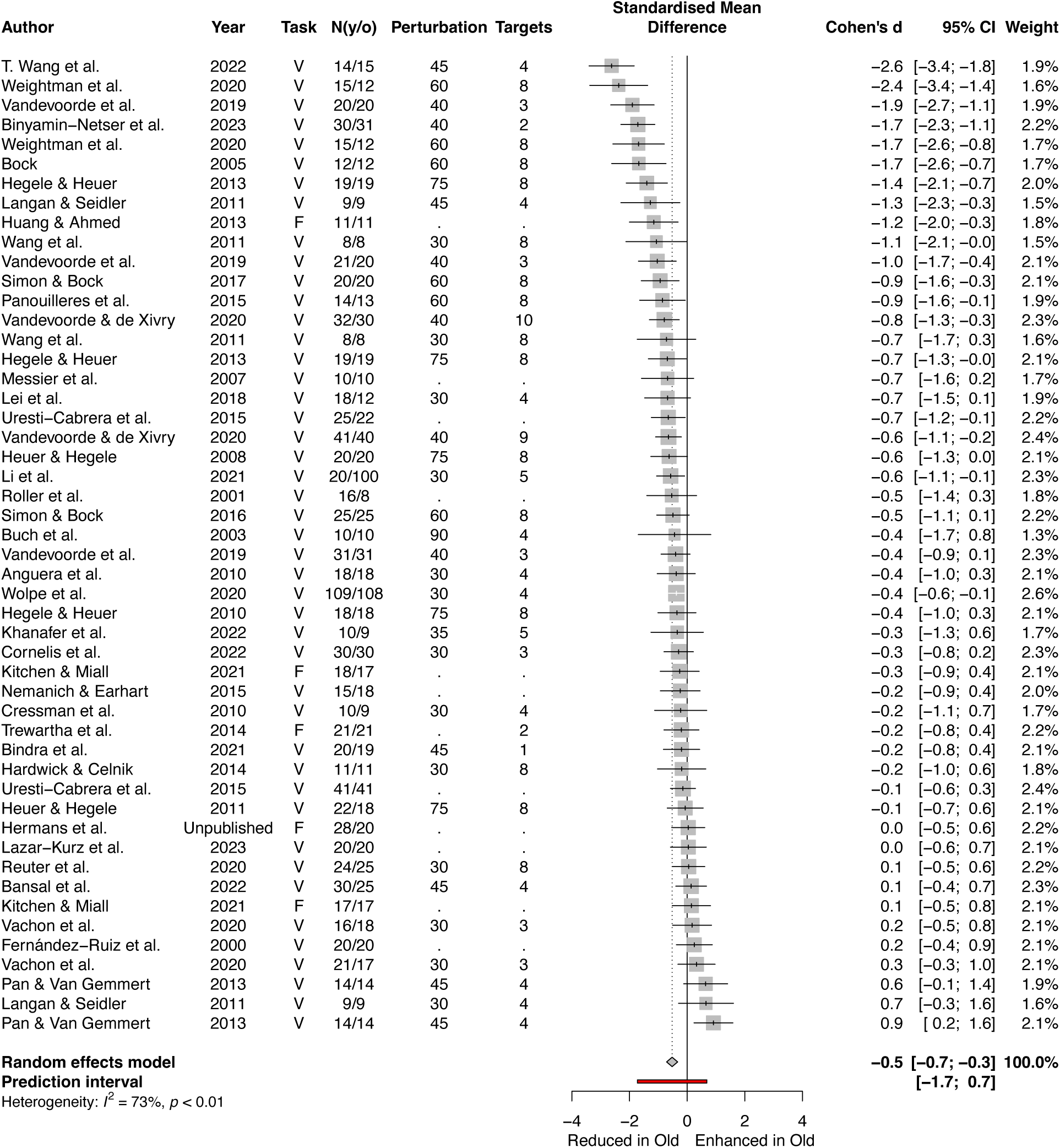
Age-related reduction in late adaptation. The meta-analysis of late adaptation included 50 datasets across 40 studies. Diamond denotes the overall effect size (Cohen’s d) and 95% confidence interval. In the Task column, V denotes visuomotor adaptation and F denotes force-field adaptation. The N column shows the number of younger (y) and older (o) participants. The Perturbation column denotes the perturbation size (°) provided in visuomotor adaptation tasks. The Targets column denotes the number of visual targets provided in the task.

#### Aging enhances implicit recalibration but reduces explicit strategy

Does this age-related decline in motor adaptation originate from a decline in implicit recalibration and/or explicit strategy? To answer this question, we focused on the 40 datasets that measured the aftereffect in the post-learning phase, providing a more targeted measure of implicit recalibration (Figure 2E). As with late adaptation, there was considerable heterogeneity across studies on the impact of aging on the aftereffect measure (I^2^ = 69%, p < 0.001), with effect sizes ranging between -1.4 and 3.1. Surprisingly, in the aggregate data, the aftereffect was significantly *enhanced* in older compared to younger adults (Figure 2F & Figure 4; d = 0.4; 95% CI [0.2, 0.5]; t(39) = 3.46; p < 0.001). This age-related enhancement was more pronounced when the meta-analysis was restricted to those that were adequately powered (power > 0.8; d = 1.1; 95% CI [0.2, 2.1]; t(6) = 2.3; p = 0.02). This age-related enhancement did not differ between experimental tasks (Q = 0.8, p = 0.4), perturbation sizes (β = 0.006, p = 0.1), or number of targets (β = -0.006, p = 0.9).

**Figure 4:**
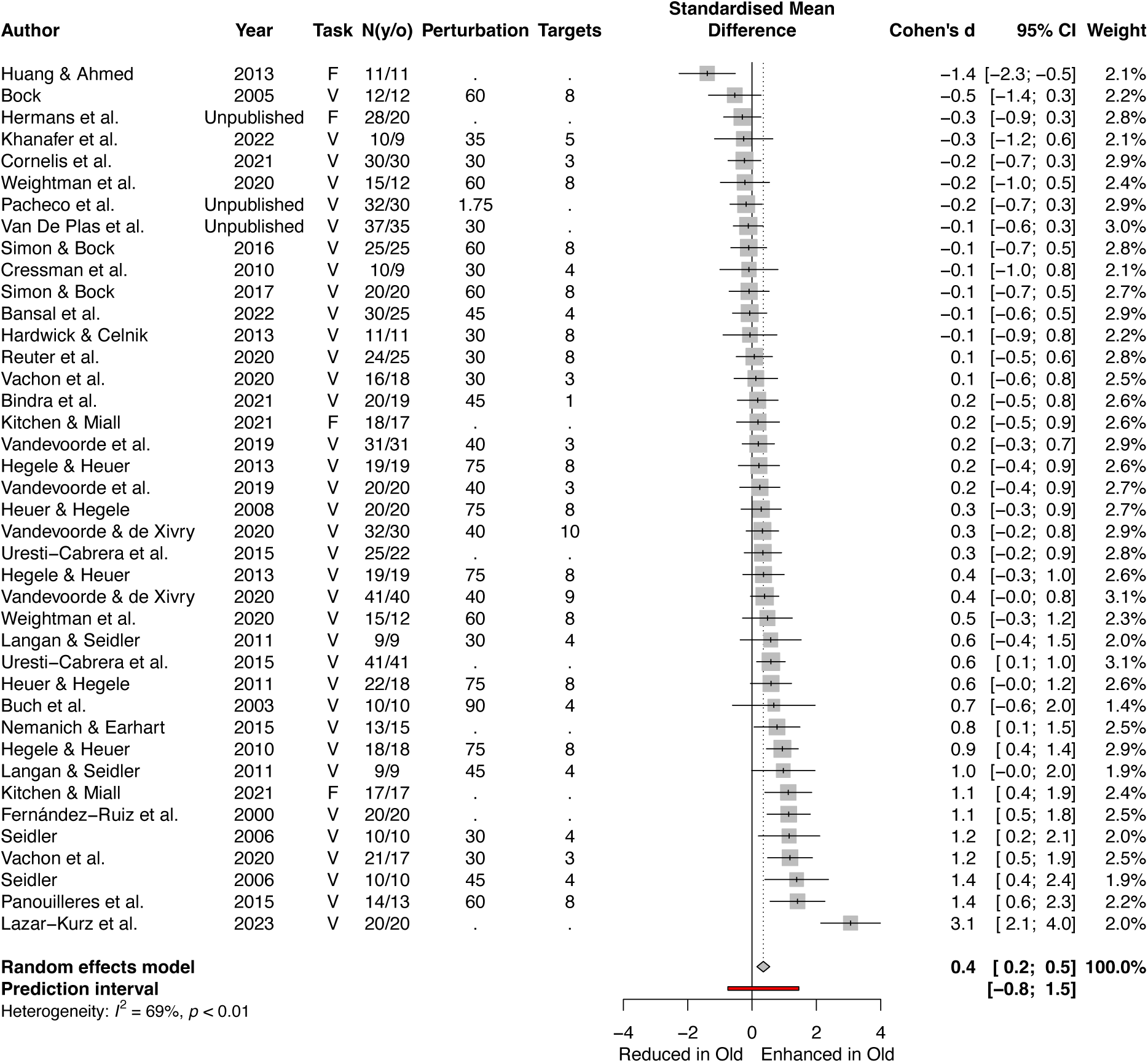
Age-related enhancement in aftereffect. This meta-analysis involves 41 datasets across 33 studies. Diamond denotes the grand effect size (Cohen’s d) and 95% confidence interval. In the Task column, V denotes visuomotor adaptation and F denotes force-field adaptation. The N column shows the number of younger (y) and older (o) participants. The Perturbation and Targets column denote the perturbation size (°) and number of target locations in visuomotor adaptation tasks, respectively.

#### Summary of meta-analysis

We can draw three conclusions from our meta-analysis. First, older participants do not perform as well as younger participants in counteracting sensorimotor perturbations. Second, there is an unappreciated age-related enhancement in implicit recalibration. Third, when considered in tandem with the age-related reduction in late adaptation, the detrimental impact of aging likely originates from an age-related decline in explicit strategy use. We test these ideas empirically in the next section.

### Experiment 1: An Age-related Reduction in Explicit Strategy

The results of the meta-analysis indicate that overall performance on sensorimotor adaptation tasks is impaired in older adults, and by inference that this deficit arises from a difficulty in identifying an appropriate aiming strategy to counteract the imposed perturbation. We directly tested this hypothesis in Experiment 1, comparing the learning performance of older and younger adults (N = 50/group) on an adaptation task in which we employed a manipulation designed to restrict learning to explicit strategy use. During the adaptation block, we imposed a 60° rotation of the cursor relative to the position of the participant’s hand. Critically, the visual feedback was provided only 800 ms after the hand reached the target distance. Delaying endpoint feedback in this manner has been shown to severely attenuate, if not outright eliminate implicit recalibration (Brudner et al., 2016; Dawidowicz et al., 2022; Kitazawa et al., 1995; Tsay, Schuck, et al., 2022). Thus, the delayed feedback task provides a method to isolate strategy-based adaptation.

Following an initial veridical feedback baseline block to familiarize participants with the task environment, the feedback cursor was rotated by 60° relative to the position of the participant’s hand (Figure 5A). To compensate for this rotation, both groups exhibited significant changes in hand angle in the opposite direction of the rotation, drawing the cursor closer to the target (baseline vs late adaptation, Younger, Median [IQR]: 55.3° [43.3°, 59.5°]; W(49) = 29, p < 0.001; d = -0.9; Older: 11.9° [-2.8°, 52.3°]; W(49) = 247, p < 0.001; d = -0.5). We did not observe a significant aftereffect for either group, indicating that participants who were using a re-aiming strategy were able to successfully ‘switch it off’ when instructed (baseline vs aftereffect, Younger, Median [IQR]: 0.8° [-0.8, 4.4]; t(44) = -1.2; p = 0.2; d = -0.4; Older: 0.8° [-2.6, 3.6]; t(49) = -1.0, p = 0.3; d = -0.3), with the exception of five younger adults who continued re-aiming. As such, these data are consistent with the hypothesis that adaptive changes observed in this delayed feedback task were primarily driven by strategic re-aiming rather than implicit recalibration.

**Figure 5:**
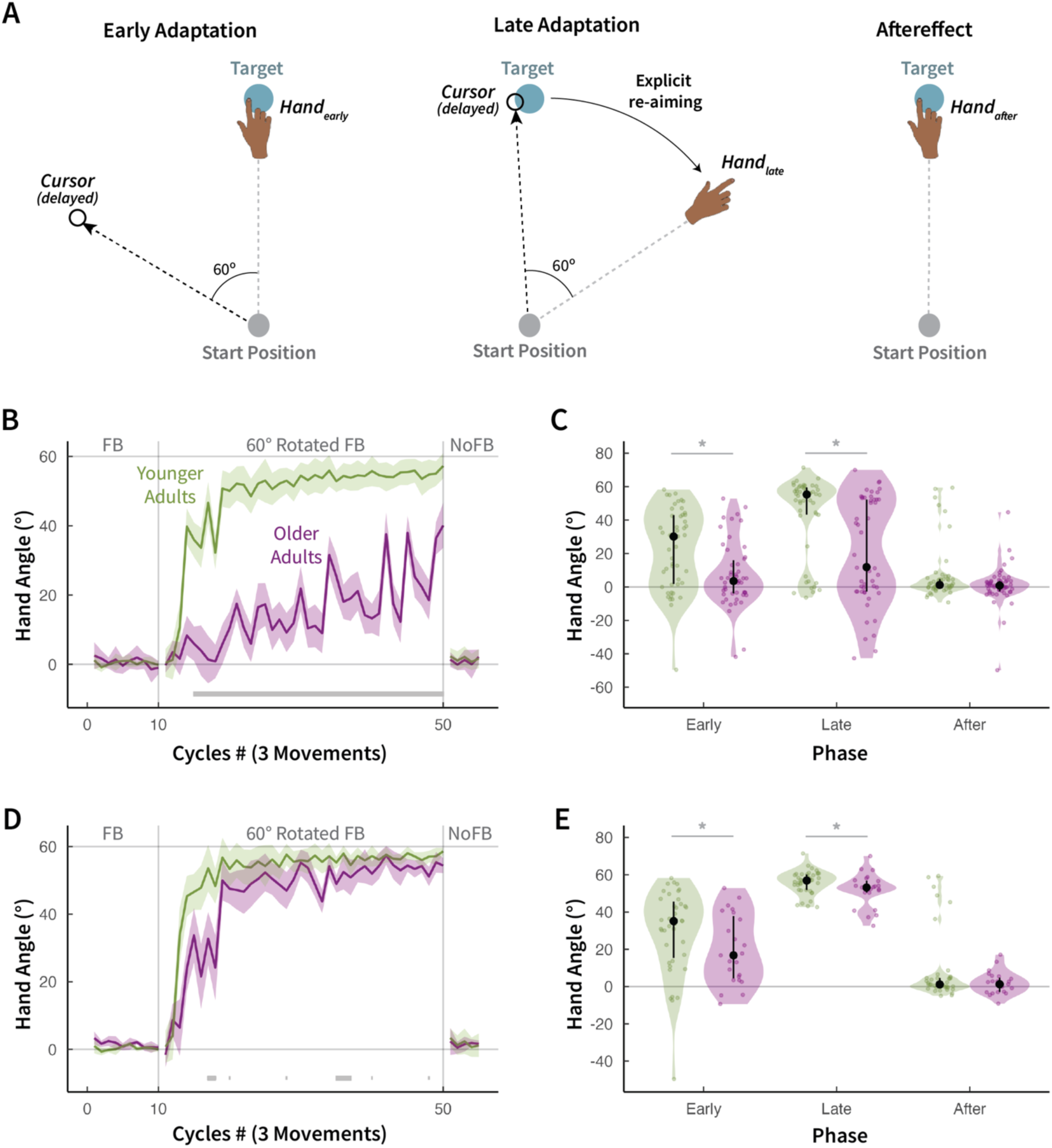
Aging reduces strategy use. **(A)** Schematic of the delayed feedback task. The 60° rotated cursor feedback (hollow black circle) was provided as endpoint feedback 800 ms after the hand reaches the target distance. **(B)** Median time courses of hand angle for younger and older adults during baseline veridical feedback (cycles 1-10), delayed feedback (cycles 11-50), and no-feedback aftereffect blocks (cycles 51-55). The shaded region denotes SEM. **(C)** Average hand angles during the early and late phases of the rotation block and the no-feedback aftereffect block. Black dot denotes the group median; error bars denote the 25% and 75% quantiles. Translucent dots represent individual participants. **(D)** Median time courses of hand angle and **(F)** mean hand angles across experimental phases for the ‘learners’. *Denotes significant difference (p < 0.05) based on t-tests between older and younger groups.

Turning to our main question, we asked how aging impacted strategy use. As depicted in Figure 5B, the learning functions were markedly reduced in older compared to younger adults throughout the perturbation block. On average, the older adult group only reached 19.8% of optimal performance (i.e., 60°) compared to 92.2% for the younger participants, pointing to a severe age-related impairment in strategy use. Based on a model-free cluster-based permutation analysis, we found a pronounced age-related reduction in hand angle during most of learning (Figure 5B gray bar: p_perm_ < 0.05, d = -0.6). These results were reinforced by a model-based analysis that showed nearly a 10-fold decline in learning rate in older compared to younger adults (learning rate: Younger, Mean [IQR] = 0.06 [0.04, 0.07]; Older, Mean [IQR] = 0.008 [0.005, 0.01]; t(52.3) = -2.7; p = 0.01, d = -0.7). Focusing on late adaptation, the median hand angle for older adults was significantly lower than for younger adults (Younger: Median [IQR] = 55.3° [43.3, 59.5]; Older: Median [IQR] = 11.9° [-2.8, 52.3]; W = 681, p < 0.001, d = -0.6). The large detrimental effect of age on late adaptation converges with the value observed in our meta-analysis.

There were no group differences in reaction time (Younger, Mean[IQR] = 675.1 ms [437.1, 806.7]; Older, Mean[IQR] = 722.7 ms [531.3, 869.6]; t(97.1) = 0.9, p = 0.4, d = 0.2). Movement time was much slower in the older adult group compared to the younger adult group (Younger, Mean[IQR] = 112.0 ms [65.2, 136.9]; Older, Mean[IQR] = 248.7 ms [79.4, 291.6]; t(57.7) = 4.1, p = 0.001, d = 1.1). While this may be a manifestation of an age-related decline in sensorimotor control, it could also reflect increased uncertainty in the older participants on their reach direction. Importantly, the explicit re-aiming deficit in the older adult group remained robust even when reaction time and movement time were included as covariates in the analyses (ANCOVA, main effect of group during the late phase of adaptation: F(1, 96) = 21.2, p < 0.001).

We then asked whether the age-related deficit in explicit strategy use arises from an inability to discover a re-aiming strategy and/or a slower rate of strategy acquisition. That is, within the timeframe of our experiment, are older adults unable to discover a successful strategy, or do they just need more practice to discover one? As shown in Figure 5C, we can clearly see that individuals varied greatly in early and late performance. This variability in both groups is not unimodal (Hartigan’s Dip Test: Younger, D = 0.07, p = 0.04; Older, D = 0.1, p < 0.001) and can be grossly binned into two subgroups: ‘Non-learners’ vs ‘Learners’. We defined non-learners as cases in which late adaptation was not significantly greater than 10% of the imposed perturbation (i.e., 6°). Based on this criterion, there were more than twice as many non-learners in the older compared to the younger group (54% vs 22%, X^2^ = 9.6, p = 0.002), indicating that the group-level strategic impairment stems in large part from a failure in strategic discovery.

The age-related failure to discover a strategy is unlikely to be due to a lack of motivation or confusion about task instructions. When we excluded participants who displayed no exploratory behavior — those with flat learning functions, defined as hand movement variance during the adaptation phase less than three times that of the baseline phase (Younger: N = 7; Older: N = 9) — we found that older adults still had more than four times as many non-learners compared to the younger group (44% vs. 9%, X^2^ = 11.3, p < 0.001). These results suggest that aging may lead to less effective sensorimotor exploration, hindering the discovery of an explicit strategy that would counteract the perturbation.

We repeated our basic analyses of the learning functions using only the data from the learners (Younger: N = 39; Older: N = 23). As depicted in Figure 5D, older learners showed subtle, but robust, deficits during the early and middle phases of learning, consistent with an age-related decline in the rate of strategy acquisition. These results were verified statistically using model-free (cluster-based permutation test: p_perm_ < 0.05, d = -0.3; Figure 5D) and model-based analyses (learning rate from an exponential model: Younger, k[IQR]= 0.1 [0.09, 0.12]; Older, k[IQR] = 0.04 [0.03, 0.04]; t(75.3) = -2.7; p = 0.009, d = 3.8). Focusing on late adaptation, the mean for the older learners was lower than for the younger learners (Younger (Median [IQR]): 56.9° [51.9, 60.2]; Older: 53.2° [50.4, 56.8]; W = 306, p = 0.04; d = -0.3. This result indicates that while both older and younger learners eventually nullify the perturbation, older learners were less efficient in discovering an effective sensorimotor strategy and were not able to nullify the perturbation to the same extent.

Taken together, Experiment 1 provides direct evidence for an age-related deficit in strategy use, corroborating the conclusions drawn from our meta-analysis of the sensorimotor learning literature.

### Experiment 2: An Age-related Enhancement in Implicit Recalibration

Unexpectedly, the results of the meta-analysis indicated that aging enhances implicit recalibration in standard adaptation tasks, although the effect size was in the low to moderate range. To follow-up on this, we compared older and younger adults (N = 50/group) in Experiment 2 on a task that isolates implicit recalibration (Figure 6A). Here the cursor follows an invariant trajectory on each trial, with its radial position matched to the participant’s hand position but the angular position shifted by a fixed angle relative to the target. Participants are instructed to ignore the non-contingent (clamped) feedback and reach straight to the target. Unbeknownst to them, a marked change in hand angle in the opposite direction of the cursor is observed, and the magnitude of the shift in late adaptation approximates that observed in a washout block recalibration (J. R. Morehead et al., 2017; Tsay et al., 2020). Although this task has been used in a previous study (Vandevoorde & Orban de Xivry, 2019), the results from different datasets even within this manuscript were mixed, leading the authors to conclude that there is no effect of aging on implicit adaptation, contradicting the findings of our meta-analysis. Here, we re-examined the impact of aging on implicit recalibration by employing a substantially larger sample size.

**Figure 6:**
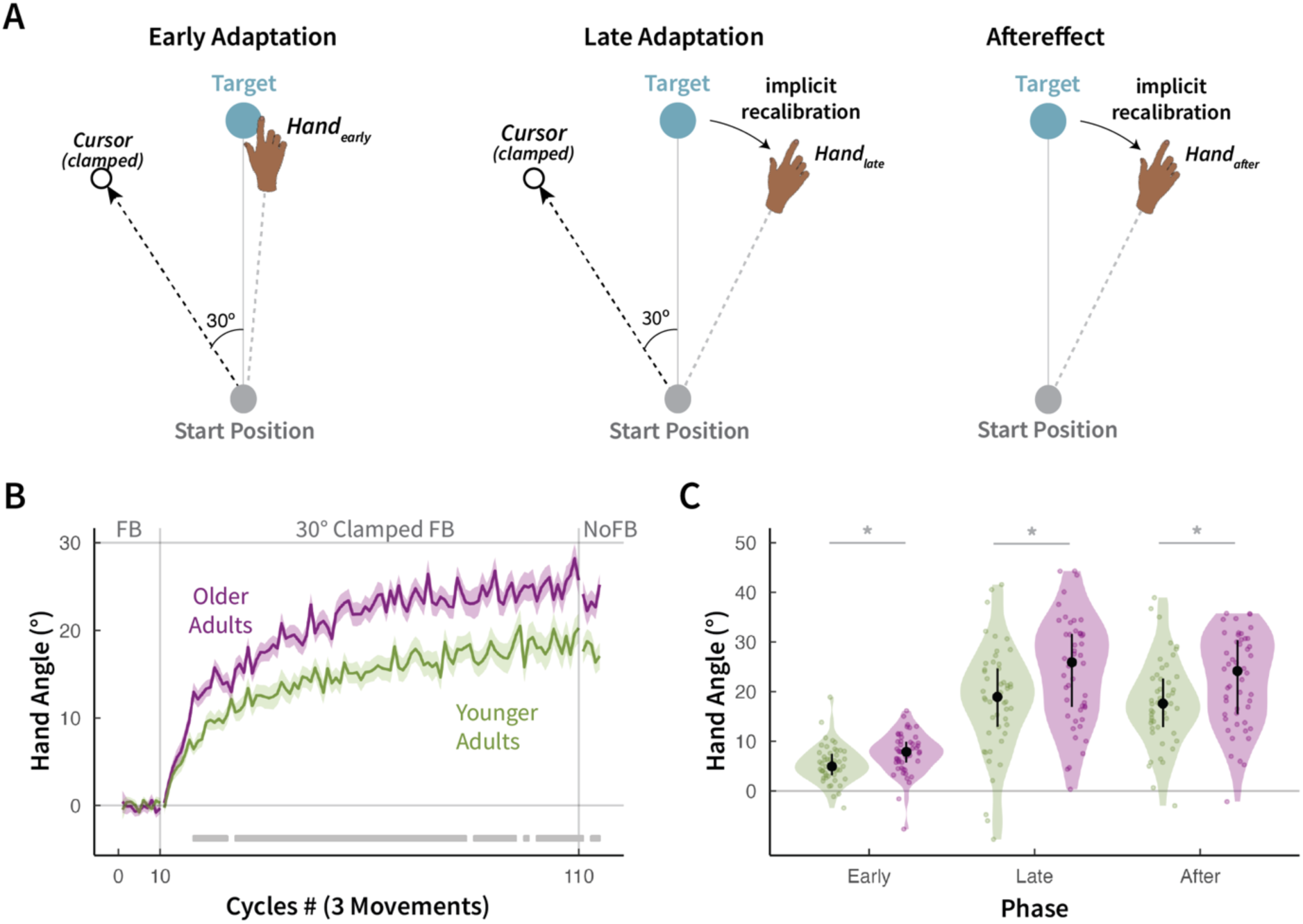
Aging enhances implicit recalibration. **(A)** Schematic of the clamped feedback task. The 30° rotated cursor feedback (hollow black circle) was provided throughout the movement. **(B)** Mean time courses of hand angle for younger and older adults during baseline veridical feedback (cycles 1-10), clamped feedback (cycles 11-100), and no-feedback aftereffect blocks (cycles 101-105). The shaded region denotes SEM. **(C)** Average hand angles during the early and late phases of the rotation block and during the no-feedback aftereffect block (black circle). Error bars represent the 25% and 75% quantiles. Translucent dots represent individual participants. *denotes significant difference (p < 0.05) based on t-tests between older and younger groups.

Once the clamped feedback was introduced, both groups exhibited a gradual change in hand angle away from the target (baseline vs late adaptation: Younger (Median [IQR]): 18.8 [12.0°, 24.6°], t(49) = -10.8, p < 0.001; Older: 25.9° [16.9°, 31.6°], t(49)= -16.2, p < 0.001). Adaptation remained robust during the no feedback aftereffect block (baseline vs aftereffect: Younger: 17.6° [12.7°, 22.6°], t(49) = 12.7, p < 0.001; Older: 24.2° [15.5°, 30.4°], t(49)= 17.0, p < 0.001). Indeed, the magnitude of the aftereffect was similar to that observed at the end of the adaptation block, consistent with the hypothesis that learning on this task is driven by implicit recalibration. Furthermore, while there were noticeable individual differences in both late adaptation and aftereffect phases, which is typical to the clamped feedback task (J. R. Morehead et al., 2017; Tsay, Irving, et al., 2023; Vandevoorde & Orban de Xivry, 2019), performance was unimodally distributed in both age groups (Hartigan’s Dip Test: Younger, D = 0.03, p = 1.0; Older, D = 0.04, p = 0.9); that is, almost all individuals exhibited robust implicit recalibration in the opposite direction of the perturbation.

Turning to our main question, we asked how aging impacts implicit recalibration. As depicted in Figure 6C, implicit recalibration was strikingly enhanced in older compared to younger adults throughout the perturbation (Younger, Mean[IQR] = 17.8° [12.0, 24.6]; Older, Mean[IQR] = 24.4° [16.9, 31.6]; t(97.3) = -3.0, p = 0.004, d = -0.6) and aftereffect blocks (Younger, Mean[IQR] = 17.2° [12.7, 22.6]; Older, Mean[IQR] = 22.3° [15.5, 30.4]; t(97.9) = -2.7, p = 0.008, d = -0.5). The medium age-related enhancement on implicit recalibration converges with the value observed our meta-analysis. Indeed, this age-related enhancement was strikingly evident in the model-free (cluster-based permutation test: p_perm_ < 0.05, d = 0.4; Figure 6B gray bar) and model-based approaches (learning rate: Younger, Mean [IQR] = 0.001 [0.0009, 0.001]; Older, Mean [IQR] = 0.001 [0.001, 0.001]; t(97.0) = 1.8; p = 0.01, d = 3.7).

Reaction times were slower in the older group compared to the younger group (Younger, Mean[IQR] = 407.1 ms [351.8, 458.5]; Older, Mean[IQR] = 471.6 ms [381.7, 517.1]; t(97.5) = 3.3; p = 0.001; d = 0.7). The older group was also slower to complete their reaches although the difference was not significant (Younger, Mean[IQR] = 115.4 ms [71.0, 127.8]; Older, Mean[IQR] = 151.5 ms [95.4, 155.8]; t(96.6) = 1.7; p = 0.09; d = 0.3). The age-related enhancement in implicit recalibration remained robust even when reaction time and movement time were included as covariates in the analyses (ANCOVA, main effect of group: F(1, 96) = 5.7, p = 0.02).

Taken together, Experiment 2 provides direct evidence for an age-related enhancement in implicit recalibration, corroborating the conclusions drawn from our meta-analysis of the sensorimotor learning literature.

## Discussion

While it is well known that aging impairs motor control, understanding the effect of aging on motor adaptation has been problematic. Previous results have been mixed, potentially due to studies often being underpowered and/or issues associated with using tasks that conflate implicit and explicit learning processes. As such, the effect of aging on different learning processes is hard to interpret, not only because the contributions of these processes are not directly measured, but also because of potential interactions between the two processes (Albert et al., 2022; Day et al., 2016; McDougle et al., 2017; Miyamoto et al., 2020; Shmuelof et al., 2012) (see (Therrien & Wong, 2022; Tsay, Kim, et al., 2023) for reviews on this topic).

We used a two-pronged approach to revisit this question. First, we conducted a meta-analysis entailing 44 motor adaptation studies that spanned three decades of work and included 2326 participants. Second, we conducted two well-powered experiments, using manipulations designed to segregate implicit and explicit processes. Convergently, the meta-analysis and empirical findings revealed an overall detrimental impact of aging on motor adaptation. With all of the caveats that go with null results, the meta-analysis indicates that the poorer performance of the older participants was observed across experimental tasks, perturbation magnitude, and task complexity (e.g., number of visual targets), highlighting the generalizability of this result.

As highlighted in the results of Experiment 1, the detrimental effect of aging on adaptation tasks stems from a decline in task-appropriate strategy use (also see: (Vandevoorde & Orban de Xivry, 2020; Wolpe et al., 2020)). There may be several, non-mutually exclusive reasons for this deficit: First, the decline in strategy could be a manifestation of a generic age-related decline in executive function (Salthouse, 2009). For example, older participants may have been less attentive to the task instructions. However, we note that even when considering participants who clearly attempted to explore different sensorimotor solutions, the rate of non-learners was much higher in the older adult group compared to the younger adult group.

Second, the age-related decline in strategy use could arise from a reduction in working memory capability. In the context of the current task, item-based working memory might be required to remember and recall the appropriate action-outcome associations for the three target locations used in Experiment 1 (McDougle & Taylor, 2019; Pellizzer & Georgopoulos, 1993). Indeed, deficits in explicit strategy use have been linked to worsened item-based working memory, suggesting that this, in part, could result in the observed deficit in older adults (Vandevoorde & Orban de Xivry, 2020).

Third, the age-related decline in strategy may be due to deficits in spatial reasoning (Anguera et al., 2010; Benson et al., 2011; Guo & Song, 2023; McDougle & Taylor, 2019; Pellizzer & Georgopoulos, 1993; Tsay, Kim, et al., 2023). Rather than learn specific associations, success on the visuomotor rotation task could come about by recognizing the spatial relationship between the cursor and movement. Recognition of this relationship would allow the participant to derive the appropriate strategy to counteract the perturbation, an algorithm that could be applied at any location. Numerous studies have demonstrated strong correlations between spatial reasoning and motor adaptation abilities in younger adults (Anguera et al., 2011; Langan & Seidler, 2011). Whether a decline in spatial reasoning causes age-related strategic changes remains to be tested.

The most striking finding in both the meta-analysis and Experiment 2 is the age-related enhancement of implicit recalibration. The absence of a deficit on this process jives with other reports of preserved implicit learning in older adults seen in other domains including reinforcement learning (Rmus et al., 2023), sequence learning (Rieckmann & Bäckman, 2009), and implicit priming (Mitchell et al., 1990). The preservation of function has been hypothesized to reflect slower age-related decline of function in subcortical areas such as the cerebellum relative to areas such as prefrontal cortex that are associated with explicit learning and cognitive control (Hedden & Gabrieli, 2005).

We are unaware of evidence indicating enhancement with age of other implicit processes. Why might this be for implicit recalibration? One hypothesis is that there is an age-related increase *in error sensitivity*: A lifetime of exposure to motor errors in older adults may have increased their sensorimotor system’s responsiveness to error. While plausible, we think this hypothesis is unlikely. Previous findings suggest that error processing is reduced, rather than enhanced, in older adults (Colino et al., 2017; Harty et al., 2017).

Alternatively, the age-related enhancement in implicit recalibration may not reflect an improvement in motor adaptation per se, but instead a change in the quality of the information and operation of processes that produce adaptation. We have proposed that implicit recalibration is driven by the mismatch between the perceived and desired position of the hand (Tsay, Kim, et al., 2022). This perceived hand position is an integration of multiple sensory inputs (e.g., proprioception for actual hand position, vision for the position of the visual cursor) and efferent inputs (e.g., predictions about the sensory outcome). In a visuomotor rotation task, the perceived hand position is biased towards the rotated visual cursor, resulting in an error that drives implicit recalibration. With an age-related decline in proprioception (Goble et al., 2009; Haaland et al., 1993; Seidler-Dobrin & Stelmach, 1998) (but see: (Cressman et al., 2010; Saenen et al., 2023)), aging could result in an enhancement of the relative weight given to the visual cursor. This would result in a greater shift of the perceived hand position toward the visual cursor and, consequently, enhanced implicit recalibration to offset this error. To test this hypothesis, or more generally explore the source of enhanced recalibration, future studies should directly assess age-related changes in multisensory integration, and how these variables relate to implicit recalibration.

In summary, our meta-analysis and empirical results jointly revealed a striking differential effect of aging on sensorimotor adaptation: Aging reduces explicit strategy use and enhances implicit recalibration. These findings deepen our understanding of the impact of aging on sensorimotor learning.

## Methods

### Meta-analysis

#### Study identification and screening

We conducted a meta-analysis to examine the impact of aging on sensorimotor adaptation. At each stage, we adhered to the Preferred Reporting Items for Systematic Reviews and Meta-Analyses (PRISMA) guidelines (Figure 1). We identified articles through a systematic search across several large research databases, including PubMed, ProQuest, Google Scholar, and PsychInfo. Our search terms included “aging,” “visuomotor rotation,” “force-field adaptation,” “prism adaptation,” and “motor learning.” We also procured articles from social media and personal correspondences with other researchers.

#### Study eligibility

We had four eligibility criteria: (1) The study involved an upper-extremity motor adaptation task in which the perturbation was created by a visuomotor displacement of the feedback or through the application of a force field. (2) The study included data comparing performance between older and younger adults. (3) The study included performance measures during the perturbation block and/or during the post-learning phase when the perturbation was removed. (4) The reports were written in English.

The following considerations were also taken into account. 1) In cases where a study featured two experiments with the same participants, we selected only the first experiment. 2) When there were multiple perturbation blocks, we extracted data from the initial round of adaptation to minimize possible session order effects (e.g., savings and/or interference (Avraham et al., 2021; Haith et al., 2015; Lerner et al., 2020; R. Morehead et al., 2015; Yin & Wei, 2020)). 3) We did not include data from experiments that involved a contextual change between learning and aftereffect phases (e.g., a change in hand usage, perturbation direction, and/or perturbation size).

We conducted our database search in Dec 2022 and identified 5443 studies. Three of the authors reviewed the list and removed 492 duplicate studies and another 4866 studies based on the title and abstract. The reviewers then independently inspected the full text of the 85 remaining studies and identified those that met the eligibility criteria. Reviewers convened to resolve any disagreements regarding inclusion criteria in order to reach consensus. The review process resulted in the removal of 40 studies that either did not include an upper extremity motor adaptation task, healthy older participants, or did not report relevant outcome measures. Following our database search, we identified 16 additional studies through private communication with colleagues and/or social media, many of which had recently been posted in preprint form or remain unpublished. In the end, there were 56 datasets from 45 studies that met the eligibility criteria.

#### Motor adaptation tasks

Our meta-analysis included two types of motor adaptation tasks that differed in the nature of the perturbation. In visuomotor perturbation tasks, participants reach to a visual target and receive visual feedback that is perturbed (e.g., rotated or translated) with respect to the position of the hand (50 datasets; Figure 2D, left). Over the course of learning, participants nullify this perturbation by moving the limb in the opposite direction of the perturbation, drawing the perturbed visual feedback closer to the target. During a subsequent no-feedback post-learning block, the perturbation and feedback are removed, and participants are instructed to abandon any explicit strategies. Despite these instructions, participants often exhibit a persistent change in heading angle in the direction of learning. This change is referred to as the “aftereffect”.

In force-field perturbation tasks, participants make a goal-directed reach to a visual target while holding a robotic device with feedback again restricted to a cursor indicating the participant’s hand position (5 datasets; Figure 2D, right). During the perturbation phase, the robot arm applies a velocity-dependent force that displaces the participant’s hand away from the target, resulting in a curved hand trajectory. Over the course of learning, participants learn to exert an equal and opposite force to counteract the force-field perturbation, with the end result being movement trajectories that are relatively straight from the start position to the target. During a subsequent post-learning block, the perturbation and feedback are removed and participants again tend to exhibit an aftereffect, now expressed as moments displaced to the opposite side of that observed in the early phases of adaptation.

#### Primary outcome measures

Our analyses focused on two primary outcome measures: First, late adaptation indexes the adaptive changes in behavior measured during the late phase of the perturbation block. Late adaptation reflects the total amount of learning arising from implicit and explicit processes. Second, the aftereffect when measured following the removal of the perturbation and feedback, provides a more targeted measure of implicit recalibration. If a measure of late adaptation or aftereffect performance was defined in the study, we used this measure in our analysis. Otherwise, we defined these measures as, the last datapoint (trial or cycle) provided in the time series data for late adaptation; the first datapoint provided for aftereffect.

#### Effect size

The meta-analysis was performed using open-source R software (Packages: *metafor* (Viechtbauer, 2015); *meta* (Schwarzer, 2007)). Cohen’s d was our standardized measure of effect size comparing younger and older groups. If the effect size was not reported, we calculated it using the F-statistic, t-statistic, or from the raw data extracted using WebPlotDigitizer (https://automeris.io/WebPlotDigitizer/). The overall effect size was computed using a random effects model. Values of 0.2, 0.5 and 0.8 for Cohen’s d represent the criteria for small, moderate, and large effect sizes, respectively. I^2^ ranges from 0 to 100%, with 25% representing low, 50% moderate and 75% high heterogeneity (Higgins & Thompson, 2002).

#### Moderator analysis and publication bias

We used Cochran’s Q test and meta-regression to examine whether the effect of aging varied between experimental tasks (visuomotor vs. force-field adaptation), the number of targets, and the size of the perturbation (limited to visuomotor rotation datasets, given that there were only 5 forcefield datasets). We used the Vevea and Hedges test to assess whether there is evidence of publication bias in the literature (Vevea & Hedges, 1995) (R package: *weightr* (Coburn & Vevea, 2019)).

### Experiments

#### Ethics statement

All participants provided informed consent in accordance with policies approved by UC Berkeley’s Instructional Review Board. Participation in the study was conducted online and in exchange for monetary compensation.

#### Participants

We recruited a total of 200 healthy adults through Prolific, an online participant recruitment platform. Of these, 100 were younger adults (age range: 19-30 years, mean ± SD age: 23.6 ± 2.7 years; 42 females), and 100 were older adults (age range: 64-79 years, mean ± SD age: 69.6 ± 4.2 years; 57 females). With the exception of two participants, all participants self-reported to be right-handed.

#### Sample size justification

Experiment 1 was designed to explore the impact of aging on explicit strategy use during sensorimotor adaptation. As such, we determined the appropriate sample size using the data from the late adaptation meta-analysis (overall effect size, mean ± SD: 0.5 ± 0.7; α = 0.05, power = 0.80, between-subject t-test). The power analysis indicated a minimal sample size of 40/group. Experiment 2 was designed to examine the impact of aging on implicit recalibration. As such, we determined the sample size using the data from the aftereffect meta-analysis (overall effect size, mean ± SD: 0.4 ± 0.6; α = 0.05, power = 0.80, between-subject t-test). The power analysis indicated a minimal sample size of 38/group. Considering that performance on online studies is more variable than lab-based studies (Tsay et al., 2024, 2021), we opted to recruit 50 participants/group for both experiments.

#### Apparatus

Participants accessed the experiment via a customized webpage using their laptop or desktop computer. They used their computer trackpad or mouse to perform the reaching task (sampling rate typically ∼60 Hz). Stimulus size and position were adjusted based on each participant’s screen size. The stimulus parameters reported below are those for a standard monitor size of 13’’ with a screen resolution of 1366 x 768 pixels.

#### General Procedure

During each trial, participants executed planar movements from the center of the workspace to a visual target. A white circle marked the center start position, and a blue circle indicated the target, both measuring 0.5 cm in diameter. On the standard monitor, the radial distance from the start position to the target was 6 cm. Targets appeared at three locations on an invisible virtual circle (30° = upper right quadrant; 150° = upper left quadrant; 270° = lower y-axis). The sequence of target locations was presented pseudo-randomly within each movement cycle (i.e., 1 movement cycle = 3 reaches: 1 reach to each target location). Movements involved joint rotations of the arm, wrist, and/or finger, depending on whether the trackpad or mouse was used. In previous validation work with this online interface and procedure, the specific movement and device used did not influence learning in visuomotor adaptation tasks (Tsay et al., 2024, 2021).

To initiate each trial, participants moved a cursor represented by a white dot (0.5 cm in diameter) to the center start position. Feedback during this initialization phase was given only when the cursor was within 2 cm of the start position. After maintaining the cursor in the start position for 500 ms, the target appeared. Participants were instructed to reach and ’slice’ through the target. There were no constraints on reaction time. To discourage mid-movement corrections, the message ‘Too Slow’ was presented on the screen for 750 ms if the movement time exceeded 500 ms. To help guide the participant back to the center start location, a veridical visual cursor appeared when the hand moved within 2 cm of the start position.

#### Experiment 1

To examine how aging impacts explicit strategy use during sensorimotor adaptation, we employed a task that isolates this process. To achieve this, we used endpoint feedback and delayed the onset of the feedback by 800 ms after the amplitude of the movement reached the target distance (Delayed Feedback Task, Figure 5A). This manipulation severely attenuates implicit recalibration (Brudner et al., 2016; Kitazawa et al., 1995); thus, changes in hand direction are almost entirely due to the deployment of a volitional aiming strategy.

There were three blocks of trials: Baseline veridical feedback (30 trials; 10 cycles), perturbed rotated feedback (120 trials; 40-cycles), and no-feedback aftereffect blocks (15 trials; 5 cycles). The baseline veridical feedback block was included to familiarize the participants with the basic reaching procedure and task requirements. On these trials, the cursor was extinguished at movement onset and only re-appeared 800 ms after the radial position of the hand reached the target distance. The position of the cursor on the veridical monitor was aligned with the position of the hand on the horizontal trackpad/mouse (as is standard when manipulating a mouse to move a cursor on a computer screen) and remained visible at that location for 50 ms. In the perturbation block, the angular position of the endpoint feedback was rotated 60° relative to the position of the participant’s hand (the angular direction was counterbalanced across participants). In the no-feedback aftereffect block, the cursor remained extinguished throughout the entire trial. The trial concluded when the target was extinguished from the screen.

Before the start of the baseline block, participants were provided the following instructions: “Please move your white cursor directly to the blue target.” Before the start of the adaptation block, participants were provided the following instructions: “Your white cursor will be offset from where you move. Hit the blue target with your white cursor.” Before the start of the aftereffect block, participants were provided the following instructions: “Your white cursor will be hidden and no longer offset from where you moved. Please move directly to the blue target.”

Through our piloting, we discovered that participants were more attentive to the task when offered a performance-based monetary bonus. Specifically, the magnitude of the monetary reward decreased exponentially with the distance between the target and the endpoint cursor position, with a maximum of $0.02 per trial if the cursor landed on the target. For example, if the cursor landed 10° away from the target (in either direction), the participant received $0.013, and if the cursor landed 40° away, they received $0.004. The cumulative monetary bonus was displayed every 15 trials. The average bonus provided per participant was $1.72.

#### Experiment 2

To examine how aging impacts implicit recalibration during sensorimotor adaptation, we compared performance of younger and older adults (N = 50/group) on a task that isolates implicit recalibration (Clamped Feedback Task; (J. R. Morehead et al., 2017). For this experiment, we used continuous feedback with the cursor remaining visible during the movement until the radial position of the hand reached the target distance. At this point, the position of the cursor was frozen and remained visible for an additional 50 ms.

There were three blocks: baseline veridical feedback (30 trials; 10 cycles), perturbed clamped feedback (300 trials; 100 cycles), and no-feedback aftereffect blocks (15 trials; 5 cycles). In the baseline veridical feedback block, the feedback cursor was aligned with the participant’s hand position, allowing the participants to become familiar with the basic reaching procedure and task requirements. In the perturbation block, we used clamped non-contingent feedback in which the radial position of the cursor was veridical but the angular position followed an invariant trajectory, displaced by 30° relative to the target (the angular direction was counterbalanced across participants). To highlight the invariant nature of the clamped feedback, four demonstration trials were provided before the perturbation block. On all four trials, the target appeared straight down (270° position), and the participant was told to reach directly to the target; however, in each case the cursor was clamped at a different offset and direction from the target. In this way, the participants could see that the spatial trajectory of the cursor was unrelated to their own reach direction. In the no-feedback aftereffect block, the visuomotor rotation was removed, and feedback was not provided.

Prior to the baseline block, participants were provided with the following instructions: “Once you see the blue target, quickly and accurately swipe towards it.” Before the adaptation block, participants were provided with the following instructions: “Your white cursor will no longer be under your control. Ignore your white cursor as best as you can and continue aiming directly towards the target.” Before the aftereffect block, participants were provided with the following instructions: “Your white cursor will be hidden and no longer be offset from where you move. Swipe directly towards the blue target.”

#### General Data Analysis

All data and statistical analyses were performed in R. The primary dependent variable was the endpoint hand angle on each trial, defined as the angle of the hand at the moment when the movement amplitude reached a 6-cm radial distance from the start position. Reaction time was defined as the interval between target onset and movement initiation, with movement initiation defined as the timepoint when the hand movement exceeded 1 cm. Movement time was defined as the time interval between movement initiation and movement completion, with the latter defined as the time at which the radial position of the hand exceeded the 6 cm target radius.

Exclusion criteria varied between experiments due to considerations regarding the learning process elicited. For Experiment 1, we excluded trials in which movement time was > 2 s; these almost always occurred on trials in which the participant failed to move the required distance. This resulted in the exclusion of 0.6% [0.0%, 3.0%] (Median [IQR]) of the trials from the younger adult group and 3.3% [1.2%, 10.0%] of the trials in the older adult group. We did not exclude reaches based on hand angle since atypical movements in this experiment might be due to trial-and-error exploration as the participants seek to identify a successful aiming strategy.

For Experiment 2, we excluded trials in which the hand angle deviated from a 5-trial trendline by more than 3 standard deviations. These datapoints are likely to reflect attentional lapses, since participants were instructed to reach directly to the target. This resulted in the exclusion of 1.4% [0.8%, 2.0%] (Median [IQR]) of the trials of the younger adult group and 1.7% [1.2%, 2.5%] of the trials of the older adult group.

We predefined three a priori learning phases: early adaptation, late adaptation, and aftereffect. Early adaptation was defined as the average hand angle over the initial ten movement cycles following the introduction of rotation block (Both experiments: cycles 11-20). Late adaptation was defined as the average hand angle over the final ten movement cycles of the rotation blocks (Experiment 1: cycles 41-50; Experiments 2: cycles 101-110). The aftereffect was defined as the average hand angle across all movement cycles during the no-feedback block (Experiment 1: cycles 51-60; Experiment 2: cycles 111-120).

#### Cluster-based Permutation Test

We employed a model-free cluster-based permutation test to identify clusters of cycles in which the hand angle differed between the two age groups (Breska & Ivry, 2019; Tsay et al., 2020; Wang et al., 2022). Wilcoxon or t-tests were used to compare each cycle of the perturbation block. Consecutive cycles with a significant difference (p < 0.05) formed a “cluster.” For each cluster, we summed W- or t-values to obtain a W/t-score statistic. We then performed a permutation test to assess the probability of obtaining a cluster of consecutive cycles. We generated 10,000 permutations by shuffling the condition labels. For each permuted dataset, we applied the same cluster identification procedure as with the actual data and computed the W or t-score statistic for each cluster. In instances where multiple clusters were found, we recorded the W or t-score statistic of the cluster with the largest value, serving as a conservative control for multiple comparisons. Clusters in the original data whose W- or t-score exceeded 95% of the W- or t-scores in the null distribution were considered statistically significant. We have included mean standard measures of effect size (Cohen’s d) of the cluster with the smallest W/t-score.

#### Learning rate Analysis

Learning rate was computed by fitting an exponential function to the data during the perturbation block of Experiments 1 and 2 (R function: drm; the ‘fct’ parameter was set to ’DRC.expoDecay’). To provide a more robust estimate of learning rate, we used a bootstrap approach by resampling the data 1000 times with replacement. Within each bootstrapped sample, we calculated the median hand angle of each trial across participants and then extracted the learning rate of the exponential function.

## Data availability

All raw data and code are available on OSF (https://osf.io/s7h5e/).

## Acknowledgements

This project was supported by the National Institute of Health (R35NS116883, R01DC0170941) and the Human Frontier Science Program (awarded to JST). The funders had no role in study design, data collection and analysis, decision to publish or preparation of the manuscript. We thank Sabrina Abram for her helpful comments on the manuscript.

## Competing interests

RI is a co-founder with equity in Magnetic Tides, Inc., a biotechnology company created to develop a novel method of non-invasive brain stimulation. The other authors declare no competing interests.

## Notes

https://osf.io/s7h5e/

